# If this title is funny, will you cite me? Citation impacts of humour and other features of article titles in ecology and evolution

**DOI:** 10.1101/2022.03.18.484880

**Authors:** Stephen B. Heard, Chloe A. Cull, Easton R. White

## Abstract

Titles of scientific papers pay a key role in their discovery, and “good” titles engage and recruit readers. A particularly interesting aspect of title construction is the use of humour, but little is known about whether funny titles boost or limit readership and citation of papers. We used a panel of volunteer scorers to assess title humour for 2,439 papers in ecology and evolution, and measured associations between humour scores and subsequent citation (both self-citation and citation by others). Papers with funnier titles were cited less often, but this appears to result from a confound with paper importance. Self-citation data suggest that authors give funnier titles to papers they consider less important. After correction for this confound, papers with funny titles have significantly *higher* citation rates, suggesting that humour recruits readers. We also examined associations between citation rates and several other features of titles. Inclusion of acronyms and taxonomic names was associated with lower citation rates, while assertive-statement phrasing and presence of colons, question marks, and political regions were associated with somewhat higher citation rates. Title length had no effect on citation. Our results suggest that scientists can use creativity with titles without having their work condemned to obscurity.

## Introduction

Do titles matter? It’s easy to find advice about constructing “good” titles for academic papers (e.g., Thomson and Kamler 2013, Silvia 2014, Saramäki 2018, Belcher 2019, Hofmann 2019, Heard 2022). By “good” titles, we generally agree that we mean those that engage readers and thus recruit them to a paper. It seems obvious that titles *should* matter in this way: they’re generally the first encounter a potential reader has with a paper, and they’re much more widely (and easily) communicated than papers themselves. Belcher (2019), for example, recommends titles that aren’t too broad, avoid abstract terms, name specific research subjects (such as species or places), include searchable keywords and verbs, and avoid cleverness or wit – among other things. There isn’t strong agreement, though, with advice from other sources sometimes concurring with Belcher’s and sometimes contradicting it. Moreover, it’s rare for advice of this sort to be supported by data.

The availability of large citation-rate datasets has made possible correlative analysis of at least one possible consequence of “good” titles: if a good title attracts readership, it should also make it more likely that the paper is cited. Conversely, papers whose bad titles repel, or at least fail to engage, readers are less likely to be cited. So what, empirically, makes a good title? The literature promises much, but delivers relatively little. For most easily-scored features of article titles, measured effects are weak (e.g., Costello et al. 2019) and inconsistent both among and within disciplines. As an example, consider title length. Most advice favours short titles, but also titles that clearly communicate an article’s contents (the fundamental contradiction between those suggestions is hard to miss). While most studies find short titles to have higher citation rates, a few have found the opposite, some find no association at all, and still others find associations that shift across disciplines (review: Heard 2021). In almost every study, though, title length explains only a small fraction of variation in citation rates. The literature for other title features (such as the use of question marks, colons, and hyphens and the inclusion of geographic place names) is similarly mixed. About the only title feature on which the literature is consistent is that titles including scientific names of genera or species are less cited than those that do not (Fox and Burns 2015, Yuret 2018, Murphy et al 2019). The picture that emerges from this work is that many features of titles are indeed associated with differences in citation rate – but that most associations are weak, and many are inconsistent. And yet it’s difficult to imagine that titles really don’t matter.

A major gap in our knowledge involves humour. Do funny titles attract reader attention, and thus increase impact? Or do they suggest that readers shouldn’t take the work seriously, and thus decrease impact? Some writing guides explicitly advise against the use of humour in titles (e.g., Thomson and Kamler 2013:85, Mack 2018:47, Belcher 2019:288). However, just three papers to our knowledge have attempted to put evidence behind this advice – likely because humour resists the kind of automated scoring that makes other features of titles easy to study. Sagi and Yechiam (2008) used panels of undergraduates to assess humour in titles of psychology papers, and found that the funniest titles were cited (slightly) less. Perhaps, they reasoned, this is because “scientific publication is considered a serious matter, and humor seems antithetical to it”. Subotic and Mukherjee (2014) attempted to replicate Sagi and Yechiam’s result (again for psychology papers), but instead found a positive effect of humour on downloads but no effect on citations. Finally, Murphy et al. (2019) found no significant effect of title humour on citation rate for ecology and entomology papers. Three other studies have examined related attributes of titles: Haslam et al. (2008) found no effect on citation of “catchiness” (a title could be catchy because it was funny, or for many other reasons) Keating et al. (2019) found a negative effect of title sarcasm, and Mammola et al. (2022) found no effect of title “pleasantness”). Together this work provides little evidence that humour helps, and yet funny titles (and the papers that bear them) are widely shared on social media and stick in memory. This incongruity suggests that humour in scientific titles deserves further study, including of the possibility that humour in titles may be correlated with other aspects of papers that influence their later citation.

We used citation rate data for 2,439 papers in ecology and evolution, taken from nine well-known journals, to ask whether humour in titles influences subsequent impact. We used self-citation data to control for possible effects of underlying differences in paper importance. We also considered two features of titles that are closely related to humour: cultural references and titles that could be considered offensive. Finally, we consider possible effects of a variety of other title features, including length, use of colons and questions, and inclusion of taxonomic and geographic names. Effects on citation rates were mostly subtle, but we present evidence that, after controlling for paper importance, funny titles increase impact. We regret, therefore, being unable to think of a funnier title for this paper.

## Methods

### Compiling papers

We compiled the titles for every paper published in 2000 and 2001 in nine well-known ecology and evolution journals: *The American Naturalist, Ecology, Evolution, Evolutionary Ecology, Journal of Animal Ecology, Journal of Ecology, Journal of Evolutionary Biology, Oikos,* and *Trends in Ecology and Evolution.* Our choice of journals was somewhat arbitrary (in particular, we did not consider impact factor), but these journals are well represented in university libraries, familiar to scientists in the field, and as a set capture both North American and European publication. Our compilation included 2,439 papers. We categorized papers as primary research articles, review articles, and “other”, with that last category including less standard forms such as “forum review” articles (*Oikos*) and “journal club” articles (*Trends in Ecology and Evolution*).

### Scoring titles

We recorded whether each paper’s title was a question or an assertive sentence (a declarative statement of a main result), and whether it was a two-part title (using a colon, dash, etc.). We also scored titles (yes/no) for the presence of acronyms or initialisms, for the inclusion of the scientific (Latin) name of a genus or species, and for the mention of a political region (country, state/province, etc.). We then assembled a group of 10 “humour scorers”, who received a spreadsheet of titles and were asked to score them for humour, offensiveness, and the presence of cultural references (allusions to books, movies, music, memes, and other non-scientific cultural knowledge). Journal names and author lists were redacted from the spreadsheets sent to humour scorers, and they were instructed not to look up any information about a paper beyond its title. Each scorer received the full set of 2,439 titles, but in a different random order. We instructed scorers to work in 20 minute sessions to avoid task fatigue, not to score more than 8 20-minute sessions in a day, and to score each title with their screen adjusted so that only that title was visible. Scorers were students or employees of the University of New Brunswick, Fredericton, New Brunswick, Canada. We had (multiple) male and female scorers and scorers originating in North America and in the Global South; their ages ranged approximately from 20 to 40. All scorers gave informed consent before their involvement, and the study was reviewed and approved by the Research Ethics Board of the University of New Brunswick (REB #2020041).

We had scorers assess humour on a 7-point scale, from zero (completely serious) to 6 (extremely funny). We did not attempt to calibrate scales across scorers. Scorers were asked to infer the author’s attempt at humour, rather than their own assessment of how funny the title was, and they were asked to ignore the subject of the article in assessing humour.

We asked scorers to identify any titles they found offensive. In contrast to the humour scoring, here we asked scorers to report their own feelings rather than their inference about the authors’ intent. Also in contrast to humour scoring, we allowed for a title to be found offensive as a result of the article’s subject (for example, a scorer might be offended by the use of humour in the title of an article addressing a very serious subject).

We asked scorers to identify titles that included cultural references of any sort (books, movies, music, memes, etc.). In a few cases, scorers reported that they suspected a cultural reference but could not identify its origin; we instructed them to include these instances. We did not restrict the age of a “cultural reference”. Thus, allusions to Vivaldi and Lil Nas X are both cultural references and are treated equally in our analyses. We acknowledge, however, that scorers might sometimes miss less current examples.

### Tracking citations

Because a minority of titles included humour or cultural references, we subset the titles database before gathering citation data. We first identified all titles for which at least one scorer recorded either a non-zero humour score or a cultural reference. There were 414 such titles, and all underwent citation tracking. From the remaining 2,025 titles, we randomly selected 650 for tracking, giving us a citation-tracked dataset of 1,064 titles. We randomized the order of titles before counting citations, because citations accumulate through time. We used Scopus^TM^ to count citations, recording the total number of citations from publication until the date of checking. We divided total citations into self- and other-citation. Self-citations were citations of the focal paper by any paper that shared at least one author; other-citations were citations of the focal paper by any paper with a non-overlapping set of authors. We use self-citations as an indicator of a paper’s intrinsic importance, reasoning that the authors’ likelihood of later citing their own paper depends on its content, not on its title.

### Data analysis

Our full data set will be available in the Supplementary Materials with the published version of this preprint. We used generalized linear models with Poisson (citation counts) or Gaussian (title attributes) error terms to explore relationships between titles and citation impact. Our primary research question was whether title humour influenced citation rate; because offensiveness and cultural references are intertwined with humour, their influences on citation was a secondary research question. We separated review articles from primary ones in these analyses, because we found authors have different practices for use of humour between article types. Finally, to complement previous studies we also examined effects on citation of several other title features, including length, use of colons and questions, and inclusion of taxonomic and geographic names. This also let us control for these variables in our analysis of title humour.

We measured agreement among humour scorers by calculating pairwise Pearson correlation coefficients among scorers and calculating Light’s (1971) kappa as an overall measure of concordance. Light’s kappa is the mean of all possible pairwise combinations of kappa scores between raters, where each κ = (*P*(*a*)−*P*(*e*))/(1−*P*(*e*)). In this expression, *P(a)* is the observed fraction of agreement and *P(e)* is the expected fraction of agreement due to chance. Kappa is often referred to as “interrater reliability”, although this implies scorers are succeeding or failing at measuring an objective underlying measurement. In our case, since humour is subjective, we are using kappa to measure agreement, not reliability, and so we avoid the latter term.

We assessed the effect of various title attributes on both total citation count and self-citation count using a series of generalized linear models, each with a Poisson error structure. Specifically, we examined the effect of the average humour, offensiveness, and cultural-reference scores for each title (Avg_humour, Avg_offense, Avg_culture), as well as article type (PrimaryReviewOther), whether the title was phrased as a question (Question), whether the title was assertive (Assertive), the presence of a colon or dash in the title (Colon), the presence of any acronyms or initialisms (Acronyms), whether the political region was noted in the title (Location), and the presence of a taxonomic name (Taxonomic_name),. For humour, we also calculated an importance-corrected citation rate as total citations divided by self citations, and tested a similar generalized linear model. We use this test primarily as a way of illustrating the importance effect, recognizing that it is not independent of the separate total- and self-citation tests. We assessed each combination of these predictor variables and ranked models according to AIC criteria – once for an analysis including all article types, and then again considering only primary research articles. We present only the best fitting model for each response variable. Because there was some multicollinearity among title characteristics, we did not include highly-correlated (>0.7) predictor variables in the same model. We did not include offensiveness or cultural-reference scores in the multivariate models as these are conceptually related to, and correlated with, humour. We examined residual plots to verify that model assumptions were met. Unless otherwise specified, for all reported results *P* < 0.01.

## Results

Citation counts for the papers we tracked were extremely variable, ranging from zero to just over 2,300 (median 64; mean 111). Unsurprisingly, review papers were cited more heavily, on average, than primary research papers; “other” papers had the lowest citation rates (Figure 1A). The citation advantage of review papers was far smaller, but still significant, for self-citation (Figure 1B). Among article types, titles from “other” papers were rated significantly more humorous than those from review and primary articles (Figure 1C). Humour did not vary significantly among journals, except that *Trends in Ecology and Evolution* (where all papers belonged to the review or “other” types) had significantly funnier titles than the rest (a higher average score and many more non-zero scores; Figure 1D). Our best fitting models and parameter estimates were similar whether we analyzed all articles or just primary research papers (compare Tables 1 and 2, for all articles, with Supplementary Materials, Tables S2 and S3, for primary research papers only). In what follows, we present only the more comprehensive analysis.

**Figure 1.**
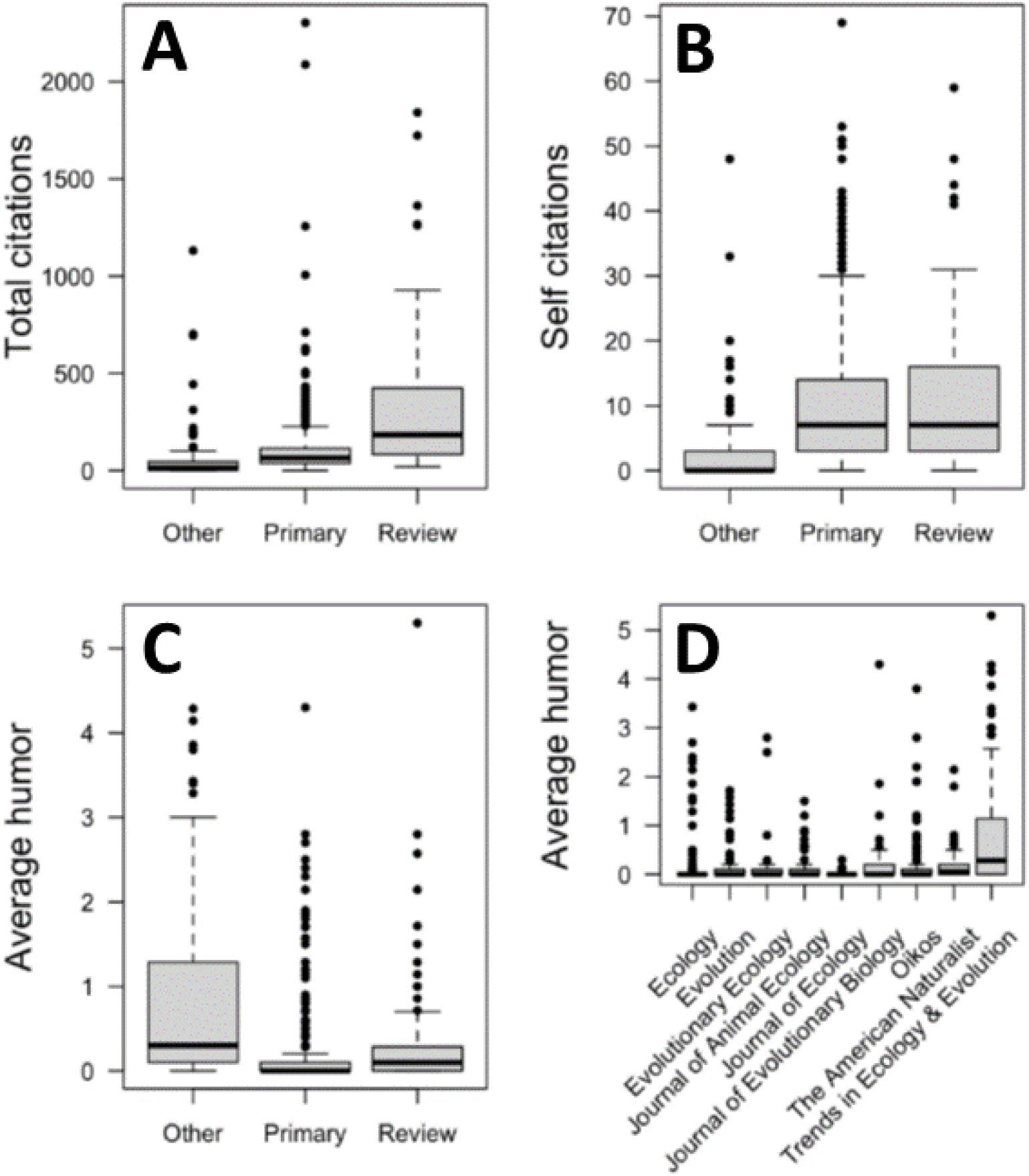
Total citations (A) and self citations (B) compared among article types (Other, Primary, or Review); and average humour scores compared among article types (C) and among journals (D). Boxplots show the median (thick horizontal line), interquartile range (25^th^ and 75^th^ percentile) for the box, and 1.5 x interquartile range for the box whiskers.

**Table 1.**
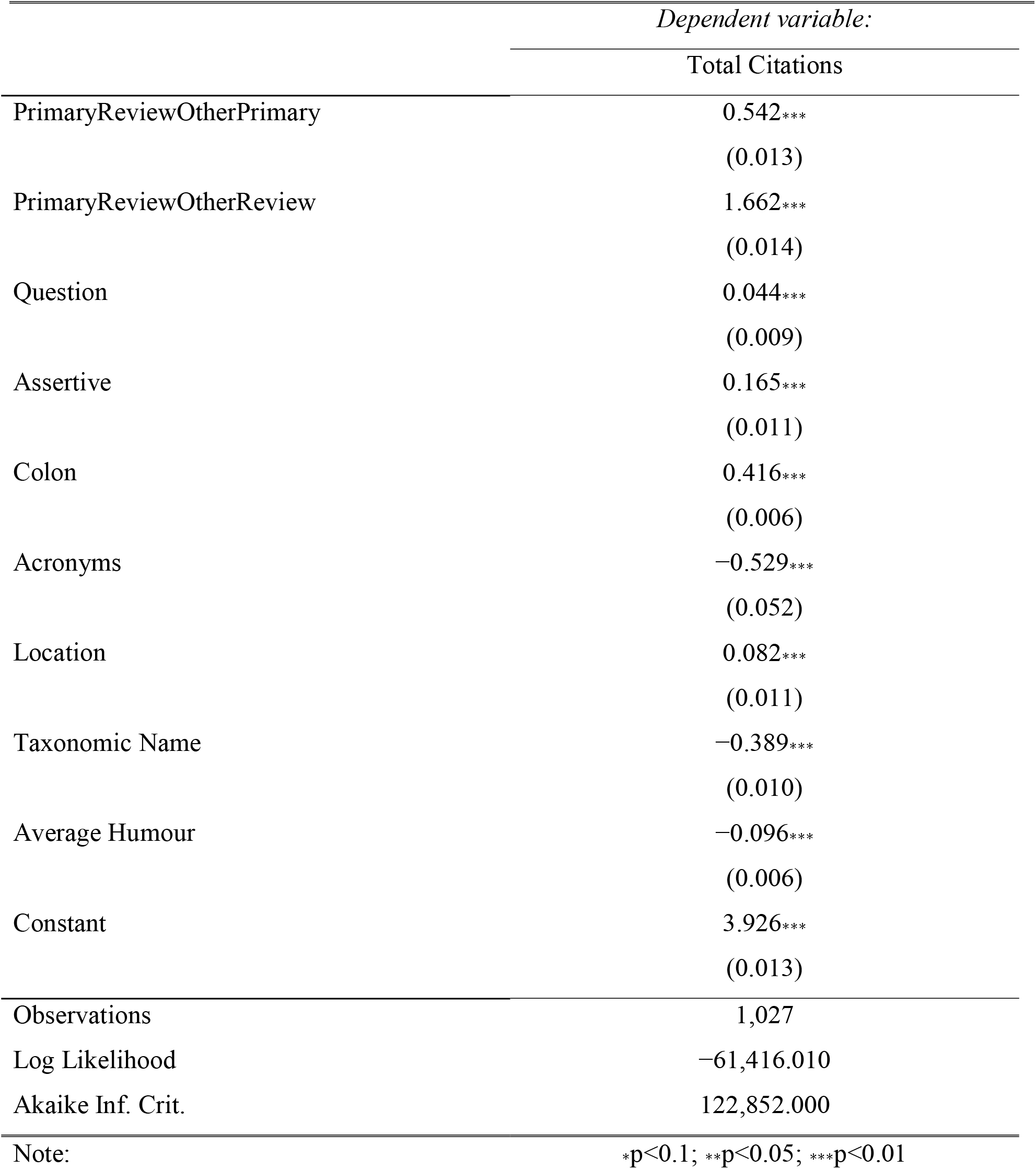
Best fitting model after AIC model selection for total citations. For each covariate, we present the log effect and (standard error) and significance level, denoted by stars.

**Table 2.**
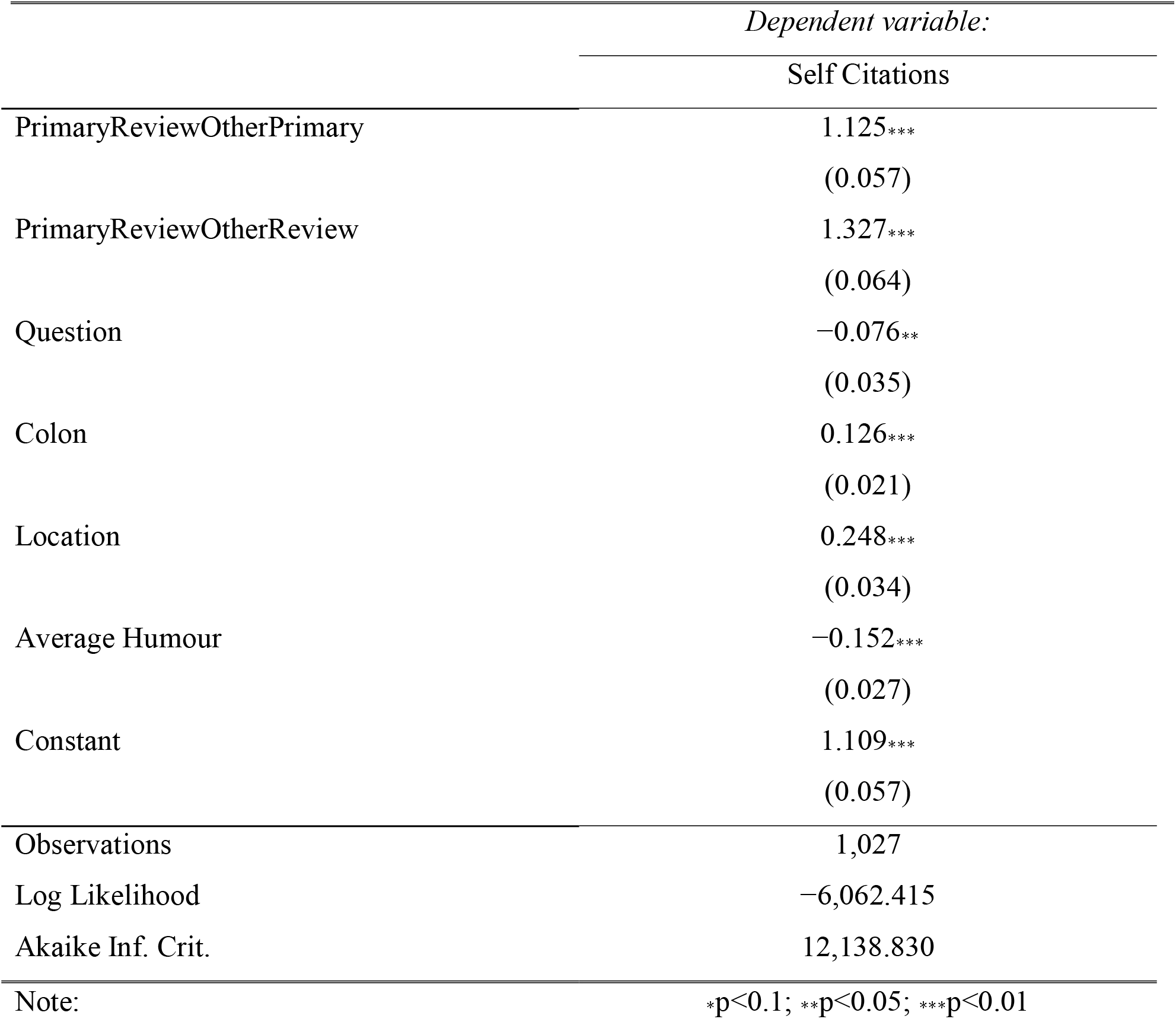
Best fitting model after AIC model selection for self-citations. For each covariate, we present the log effect and (standard error) and significance level, denoted by stars.

Few titles were funny: only 414 of 2,439 papers were assigned a non-zero humour score by even one scorer, and only 60 had at least 11 humour points (as they would if all scorers gave them the minimum non-zero humour score, or two scorers gave them the maximum score). The median humour score was zero (mean 0.096).We saw fairly low, but non-zero, agreement among scorers in their assessment of title humour. The overall concordance score (Light’s kappa) was just 0.34; most pairwise (Pearson) correlations had *r* < 0.5, and over a third had *r* < 0.35 (Figure 2, and precise correlations in Supplemental Materials, Table S4). The title with the highest humour score was “Nice snake, shame about the legs”; this title also tied for the highest offensiveness score. Other titles with relatively high humour scores included “Some Like it Hot: Intra-Population Variation in behavioral Thermoregulation in Color-Polymorphic pygmy Grasshoppers”, “Is it Time to Bury the Ecosystem Concept? (With Full Military Honors, of Course!)”, and “The Competition Colonization Trade off is Dead; Long Live the Competition Colonization Trade off”. Only the first title received a non-zero humour score from every scorer.

**Figure 2:**
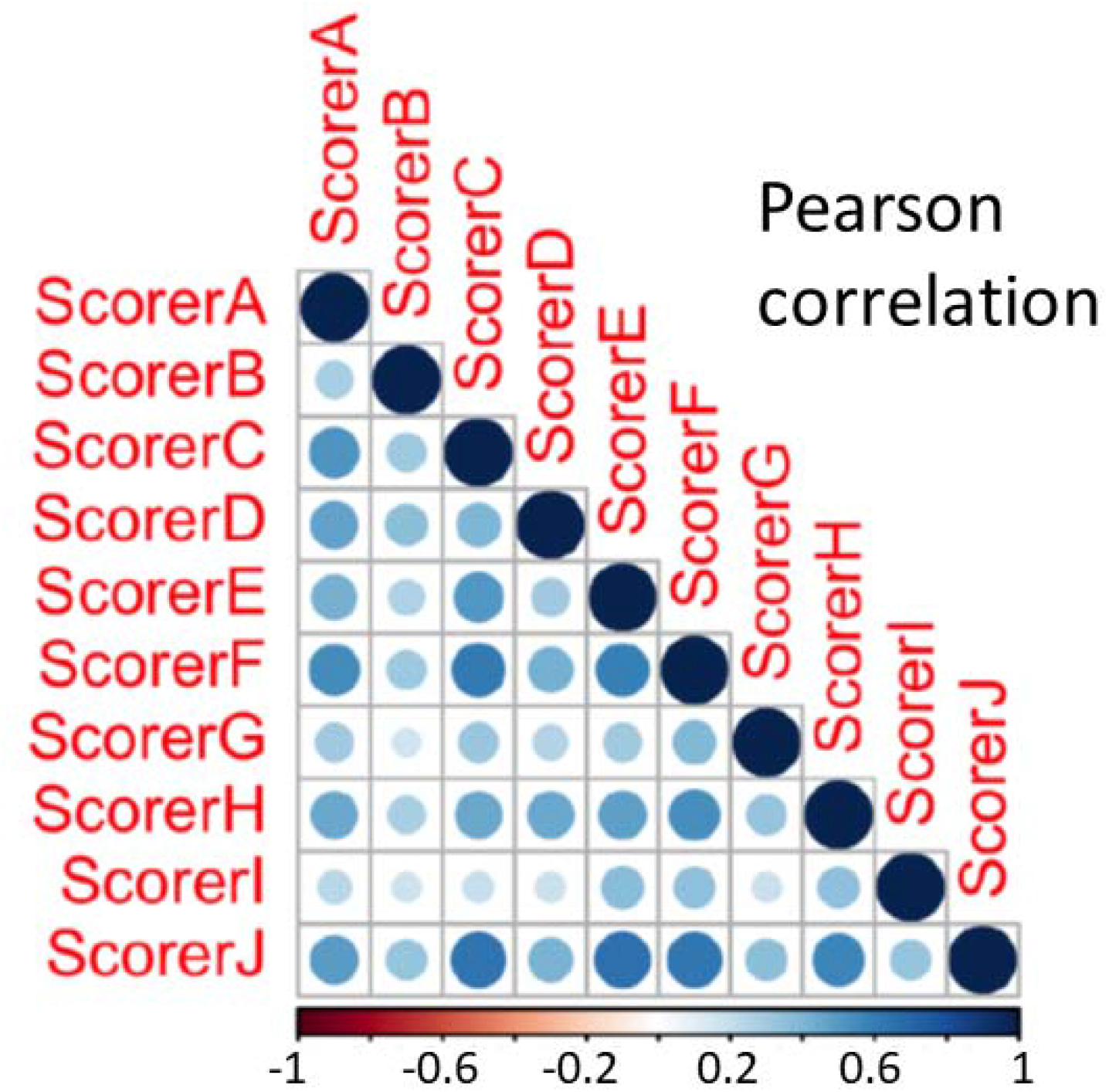
Concordance among scorers for title humour. The matrix shows Pearson correlation coefficient (*r*) for each pairwise combination of scorers, across all scored titles. The overall concordance, measured by Light’s kappa, was 0.34.

Our best-performing models (Tables 1 and 2) suggested contributions to citation rate from title humour but also from phrasing titles as questions, including colons, acronyms, locations, and taxonomic names, and (for all article types but not for primary research papers alone) phrasing titles as assertive statements. However, some of these effects were weak (see below).

After we controlled for other predictors, total citations declined with average title humour (Figure 3A). The effect was relatively small, with a decrease of 4% in total citations for each 1 point increase in average humour score, but this equates to a difference of 20.4% between the least and most humorous titles. There is, however, an important qualification: the pattern was similar, but much stronger, for self citations, with an 82% decrease for the most humorous titles (Figure 3B). Thus, after correcting for underlying paper importance, funny title are cited more, not less (Figure 3C), with a 23% increase for each 1 point increase in humour score.

**Figure 3:**
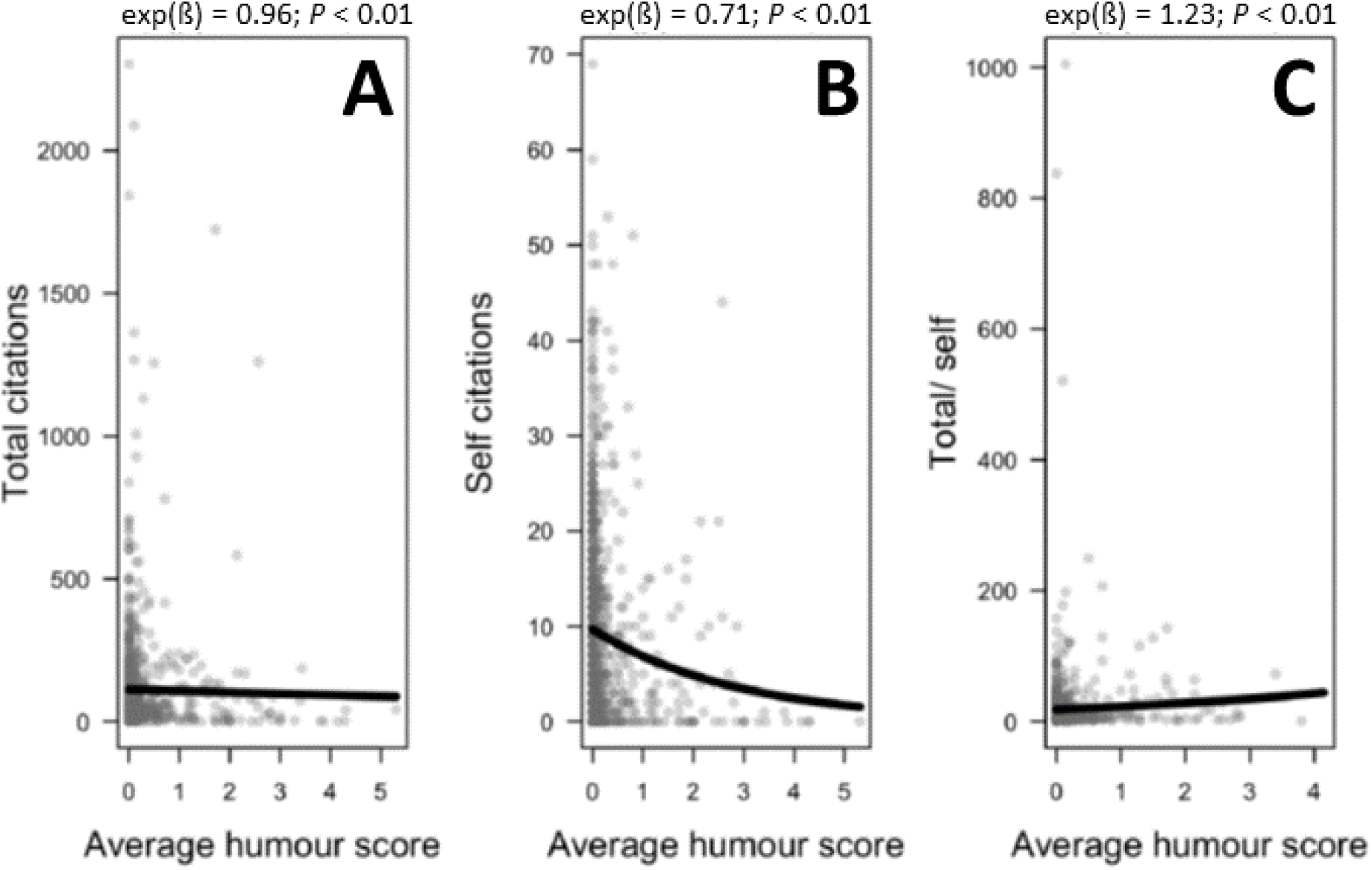
Humour and citation rates. Both total citations (A) and self citations (B) significantly decreased with higher humour scores. However, the effect size is much larger for self citations, and the ratio of total to self (C) citations *increases* with humour score.

While we did not include offensiveness or cultural references in our AIC modeling, we examined their association with citation rates in isolation. Offensive titles were rare, with only 19 of 2,439 titles scored as offensive by even a single scorer (median 0, mean 0.06). Citation rates declined with average offense score (Figure 4A). However, as for humour, there was an even stronger decline for self-citations (Figure 4B), suggesting that less important papers are given titles that our scorers judged offensive. Titles including cultural references show a pattern of increasing citation (Figure 4C), despite fewer self-citations (Figure 4D, again suggesting lower underlying paper importance). Interestingly, the detection of cultural references by our scorers was quite imperfect. 75 titles were recorded as including a cultural reference by at least one scorer, but only 5 were so recorded by a majority of scorers and none by all scorers..

**Figure 4:**
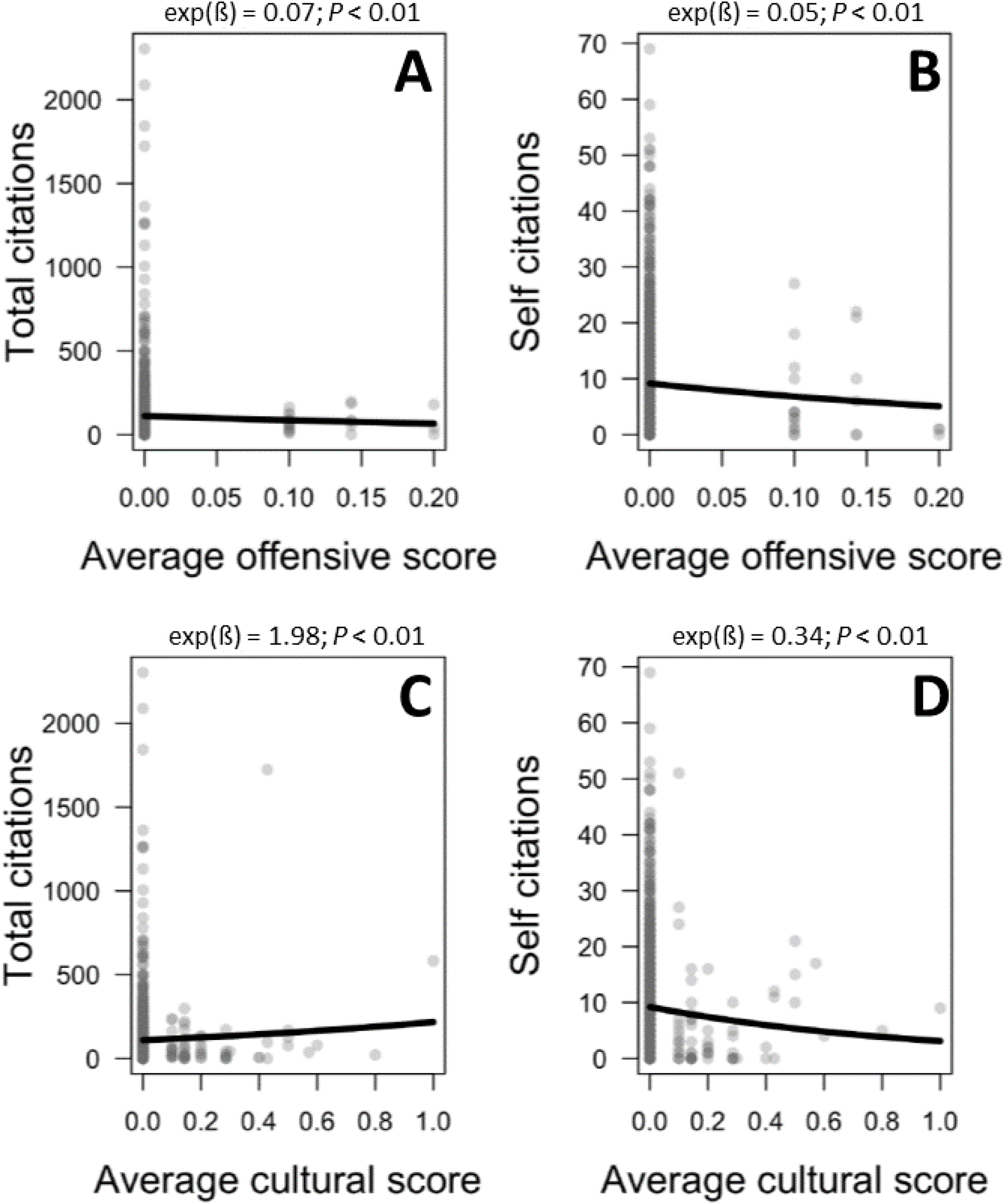
Offensive titles, cultural references, and citation rates. Total citations decreased significantly with higher offensive scores (A), but self citations decreased more strongly (B). The inclusion of cultural references was associated with higher total citations (C) but with *lower* self citations (D).

Several other characteristics of paper titles were significant predictors of citation counts in the AIC model, but most of these effects were relatively weak. Titles with colons or question marks, those phrased as assertive statements, and those including names of political regions were more highly cited (Table 1 and Supplementary Material Figure S1, upper row), although only the colon effect was strong and the “assertive statement” effect disappeared when we analyzed only primary research articles; Supplementary Material Table S2). Effects on self-citation were mostly very weak (Supplementary Material Figure S1, lower row), except that titles mentioning political regions had moderately more self-citations. Finally, title length was excluded from all AIC models (Tables 1, 2) and made little difference to either total or self citation rates viewed in isolation (Supplementary Materials Figure S2).

We found stronger effects for the inclusion in titles of acronyms and taxonomic names. Each was associated with a sharp decrease in citation rates (acronyms 41%, Figure 5A; and taxonomic names 32%, Figure 5C). These effects cannot be explained by paper importance, as the inclusion of acronyms was not associated with self-citation (Figure 5B) and the inclusion of taxonomic names was associated with slightly higher self-citation (Figure 5D).

**Figure 5:**
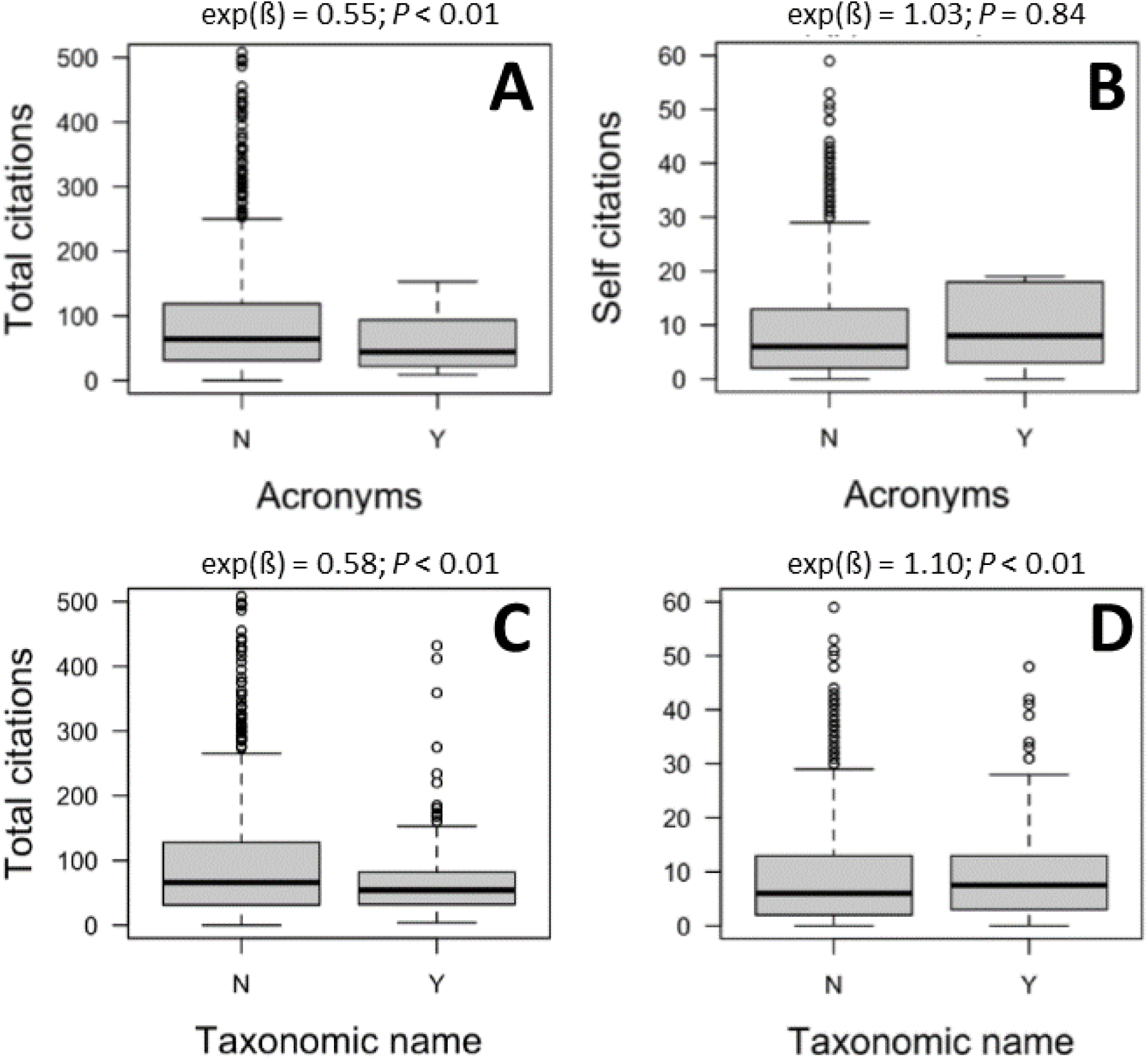
Acronyms, taxonomic names, and citation rates. The inclusion of acronyms was associated with a significant decrease in total citations (A), but was not associated with self citations (B). The inclusion of taxonomic names was associated with a strong decrease in total citations (C) but a slight increase in self citations (D). Boxplots show the median (thick horizontal line), interquartile range (25^th^ and 75^th^ percentile) for the box, and 1.5 x interquartile range for the box whiskers.

## Discussion

Despite the widespread availability of clear and firm advice on constructing “good” titles, the most striking pattern we document is simply that few easily measured attributes of titles seem to have strong associations with citation rates. This is broadly consistent with the literature (e.g., Costello et al. 2019, Murphy et al. 2019, Mammola et al. 2022; review: Heard 2021).

There were some differences in humour scores among the three article types we distinguished. In particular, “other” articles (forum review and journal club papers) had both the highest humour scores and the lowest citation rates. This can account for the higher average humour scores for one journal (*Trends in Ecology and Evolution*), where the bulk of “other” papers were published. Otherwise, though, article type didn’t drive the patterns in citation rate we observed, as analyses restricted to primary research articles had very similar results to those including all three article types.

Our analysis suggests that humour in the title can increase a paper’s impact. It is true that the simplest analysis, correlating total citations with humour score, finds a (weak) negative relationship. However, such an analysis fails to account for the possibility that authors are less likely to use humour in titling their more important papers. Our self-citation data strongly suggest that this is true: papers with funnier titles are subsequently cited less *by their own authors*. Since authors don’t need titles to alert them to their own papers, self-citation provides a title-independent estimator of importance – unlike other-citations, for which effects of title and underlying importance on citation are inextricably confounded. Because the decline in self-citation with humour score is much steeper for self-citations than for other-citations, funny titles are actually over-cited, not under-cited, after correction for paper importance (Figure 3C).

Earlier literature has not considered the possibility of confounding between title humour and paper importance. An analysis for psychology papers by Sagi and Yechiam (2008), which found a negative association between total citations and title humour, did not attempt any correction for paper importance, via self-citation or otherwise. As a result, that analysis may well have drawn precisely the wrong conclusion. The same issue applies to analyses by Subotic and Mukherjee (2014) and Murphy et al. (2019), both of which found no effect of humour on total citation but, again, did not correct for paper importance. Advice to avoid humour in paper titles (e.g., Thomson and Kamler 2013:85, Mack 2018:47, Belcher 2019:288) is thus not well founded in evidence – at least, not if the concern is citation impact.

Scientists sometimes express two related worries about the use of humour: that funny titles might be seen as offensive, and that funny titles will be misunderstood by those who don’t share the author’s cultural background. Our data suggest three things about this. First, if these things happen, they don’t affect citation much. Papers with titles identified as offensive were indeed cited less, but as for humour, analysis of self citations suggests that this can be more than explained by the use of such titles for less important papers. Second, the low concordance among our scorers suggest that even with a group of scorers of relatively homogeneous cultural background, opinions about humour and offense vary widely. The simultaneous existence of *South Park* and *The Satanic Verses* should make it obvious that both humour and offense are deeply personal, and both will sometimes be perceived even when neither is intended. Third, even though some readers will miss cultural references in titles (it was commonplace for our scorers to differ in their detection), this does not interfere with discovery or impact of the papers: the use of cultural references was strongly associated with increased citation rates.

Other features of titles are significantly associated with citation rates, but most of the effect sizes are small – as has generally been true in previous studies. Citation rates are higher for two-part titles (those with colons, dashes, etc.) and a little higher for question and assertive sentence titles. Inclusion of a geographic region name increases citation a little, consistent with some other studies (Rostami et al. 2013, Nair and Gibbert 2015, Murphy et al. 2019) but contrasting with others (Jacques and Sebire 2010, Paiva et al. 2012, Abramo et al. 2016, Alimoradi et al. 2016, Yuret 2018, Costello et al. 2019). However, analysis of self citation suggests that this is likely explained by a tendency for authors to use geographic names in their more important papers. We do not have an explanation for this tendency, which surprised us. Title length, which is one of the most frequent targets of well-meaning advice, had virtually no effect on citation. This is broadly consistent with the literature (review: Heard 2021): shorter titles are sometimes found to be cited more, and sometimes found to be cited less, but the effects vary from weak to very weak. Keeping titles short may help typesetters, but seems to have no implication for authors or readers.

There were larger effects for taxonomic names: their inclusion is associated with a steep (32%) reduction in citation. The negative effect of taxonomic names in titles is one of the few citation effects to be consistent across studies (Fox and Burns 2015, Yuret 2018, Murphy et al. 2019). Readers appear to behave as if inclusion of a taxonomic name signals narrower scope of, and thus narrower interest in, a paper. This could be a reliable signal (papers including taxonomic names may, on average, genuinely be of narrower scope) or a misperceived one (with readers being deterred from papers that really are relevant to them). Since *self*-citations don’t decline with the inclusion of a taxonomic name, we suspect that misperception is often involved. Authors may therefore wish to consider removing scientific names of taxa from titles.

Finally, we were surprised by the strong pattern for acronyms. Despite our deep familiarity with – perhaps even love for – acronyms (Barnett and Doubleday 2020), their appearance in a title is associated with a 41% decrease in citation rates, and this can’t be explained by variation in paper importance. There were already good reasons to reduce our use of acronyms in writing; their apparent effect on citation impact may add another.

There is, of course, an important assumption behind our choice of citation rate as a variable to correlate with features of titles. Citation rate is only of interest if it says something useful about the reach or impact of a paper. Given that science is a fundamentally cumulative process, and given that modern citation practices involve an ethical responsibility to cite influential work, citation rate really does seem likely to be measuring something useful. In a few cases, of course, a paper may be heavily cited because it’s wrong – for example, as an example of how an analysis can go astray – but we doubt that such citations account for a significant fraction of our database.

Ultimately, the factors that explain the citation impact of a paper are sure to be numerous, interrelated in complex fashion, and extending far beyond just the title. However, because titles are the first point of contact with a paper for most readers, we suspect interest in their construction will remain strong. In a sense, our results are mostly good news for authors: few title features (barring acronyms and taxonomic names) work against citation. That means scientists can use titles creatively, even inserting touches of humour (Heard 2014), without fear of their work ending up in undeserved obscurity.

## Acknowledgements

We thank our scorers, who read more paper titles in a couple of weeks than some people do in a career. Lyndsey Burrell grappled bravely and well with searching and summarizing a rather peculiar literature. We are grateful to Joel Dacks, an anonymous reviewer, and members of the PEER Group, Dept. of Biology, University of New Brunswick, for helpful comments on the work and the manuscript. This research was funded by the Natural Sciences and Engineering Research Council of Canada via a Discovery Grant to SBH.

## Supplementary Material

*Supplemental Tables and Figures:*

*Table S1. Full dataset used in analyses (to be provided with the published version).*

*Table S2. AIC-selected model for total citations, primary research papers only.*

*Table S3. AIC-selected model for self citations, primary research papers only.*

*Table S4. Pearson correlations among scorers for title humour.*

*Figure S1. Associations with total citation rates (top row) and self citation (bottom row) for two-part titles (“colon”), question titles, assertive-sentence titles, and titles including names of political regions.*

*Figure S2. Title length and rates of total (A) and self (B) citation.*

