## Supplemental Figure 1 for "If this title is funny, will you cite me? Citation impacts of humour and other features of article titles in ecology and evolution"

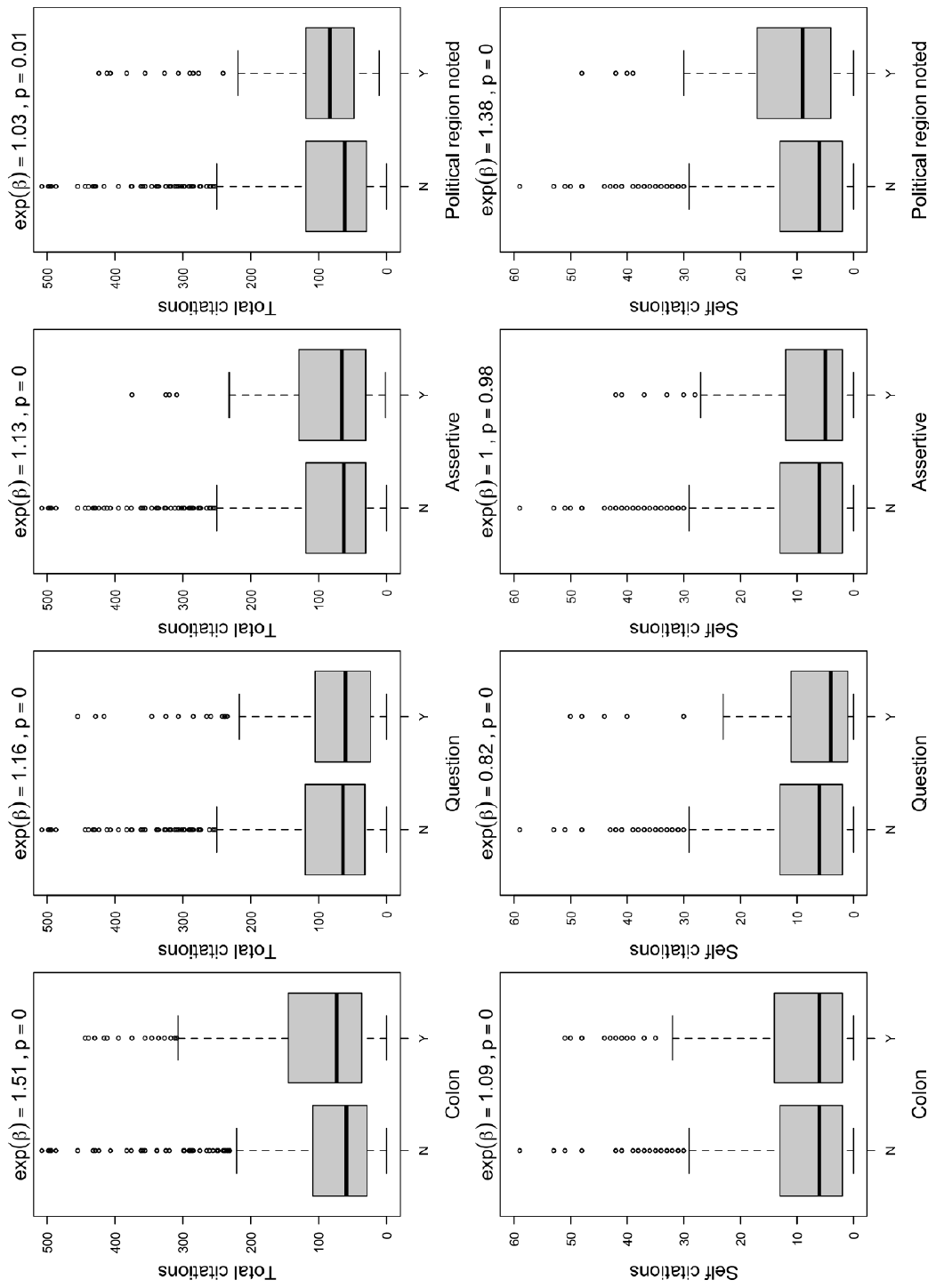

Figure S1. Associations with total citation rates (top row) and self citation (bottom row) for two-part titles
