## Supplemental Figure 2 for "If this title is funny, will you cite me? Citation impacts of humour and other features of article titles in ecology and evolution"

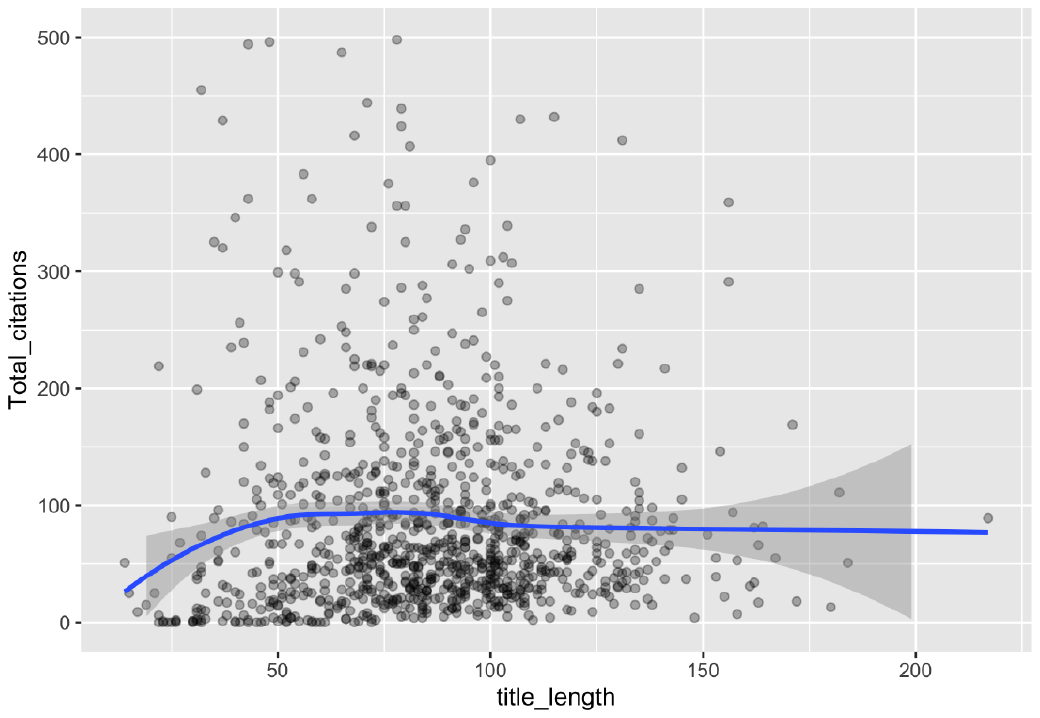

A

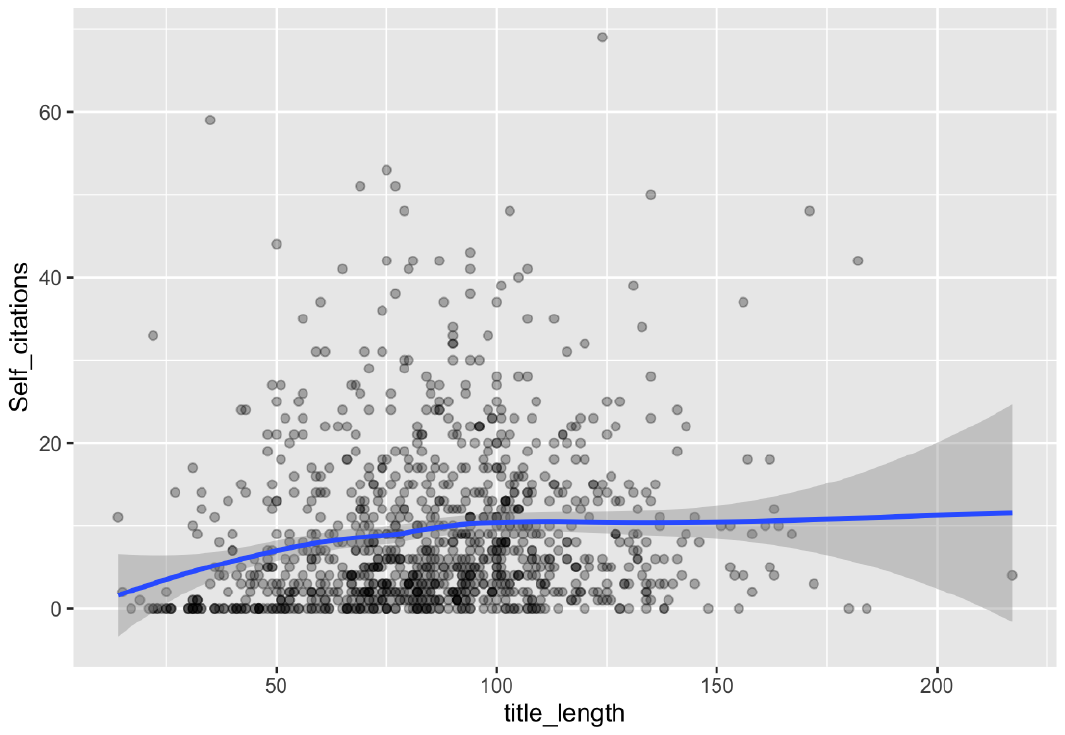

B

Figure S2. Title length and rates of total (A) and self (B) citation.
