## Supplemental Table 2 for "If this title is funny, will you cite me? Citation impacts of humour and other features of article titles in ecology and evolution"

Table S2: Best fitting model after AIC model selection for total citations for PRIMARY

articles only. For each covariate, we present the log effect and (standard error) and significance level, denoted by stars.

|  | *Dependent variable:* |
| --- | --- |
|  | Total_citations |
| QuestionY  ColonY  AcronymsY  LocationY  Taxonomic_nameY  avg_humour  Constant | 0.094∗∗∗  (0.012)  0.299∗∗∗  (0.007)  −0.517∗∗∗  (0.052)  0.205∗∗∗  (0.012)  −0.395∗∗∗  (0.010)  −0.112∗∗∗  (0.010)  4.523∗∗∗  (0.005) |
| Observations  Log Likelihood  Akaike Inf. Crit. | 819  −39,417.760  78,849.520 |
| Note: | ∗p<0.1; ∗∗p<0.05; ∗∗∗p<0.01 |
