## Supplemental Table 3 for "If this title is funny, will you cite me? Citation impacts of humour and other features of article titles in ecology and evolution"

Table S3: Best fitting model after AIC model selection for self citations for PRIMARY

articles only. For each covariate, we present the log effect and (standard error) and significance level, denoted by stars.

|  | *Dependent variable:* |
| --- | --- |
|  | Self_citations |
| ColonY  LocationY  avg_humour  Constant | 0.055∗∗  (0.023)  0.260∗∗∗  (0.035)  −0.126∗∗∗  (0.035)  2.250∗∗∗  (0.015) |
| Observations  Log Likelihood  Akaike Inf. Crit. | 819  −4,902.983  9,813.965 |
| Note: | ∗p<0.1; ∗∗p<0.05; ∗∗∗p<0.01 |
